# deepManReg: a deep manifold-regularized learning model for improving phenotype prediction from multi-modal data

**DOI:** 10.1101/2021.01.28.428715

**Authors:** Nam D. Nguyen, Jiawei Huang, Daifeng Wang

## Abstract

The biological processes from genotype to phenotype are complex involving multi-scale mechanisms. Increasing multi-modal data enables deeper understanding of underlying complex mechanisms in various phenotypes. However, integrating and interpreting such large-scale multi-modal data remains challenging, especially given highly heterogeneous, nonlinear relationships across modalities. To address this, we developed an interpretable regularized learning model, deepManReg to predict phenotypes from multi-modal data. First, deepManReg employs deep neural networks to learn cross-modal manifolds and then align multi-modal features onto a common latent space. This space aims to preserve both global consistency and local smoothness across modalities and reveal higher-order nonlinear cross-modal relationships. Second, deepManReg uses cross-modal manifolds as a feature graph to regularize the classifiers for improving phenotype predictions and also prioritizing the multi-modal features and cross-modal interactions for the phenotypes. We applied deepManReg to recent single cell multi-modal data such as Patch-seq data including transcriptomics and electrophysiology for neuronal cells in the mouse brain. We show that deepManReg significantly improves predicting cellular phenotypes and also prioritizing genes and electrophysiological features for the phenotypes. Finally, deepManReg is open-source and general for phenotype prediction from multi-modal data. deepManReg is open-source available at https://github.com/daifengwanglab/deepManReg.

## 1 Introduction

Recent large-scale multi-modal data such as various next generation sequencing data allows a deeper understanding of cellular and molecular mechanisms from genotype to phenotype. Also, many of those data have been used to predict phenotypes, transforming the bioinformatics research from descriptive to predictive [8]. However, it is still challenging to integrate and analyze those multi-modal data which are typically high-dimensional and heterogeneous across modalities for phenotype prediction. In particular, cross-modal features likely have the nonlinear relationships that many computational methods may miss for predicting phenotypes [19]. For example, feature extraction and feature selection are widely used to reduce the dimestionality for prediction. However, the unselected features also likely have useful relationships (likely nonlinear) which potentially are able to contribute to prediction [17]. Therefore, systematically identification of nonlinear features across modalities is key to improving phenotype prediction from multi-modal data. Manifold alignment is a widely used technique that simultaneously reduces the dimensions of multiple data types and preserves the geometric nonlinear local structures in and between data types (which is also known as multiview nonlinear dimensionality reduction [22, 11, 13]). However, such methods suffer from a trade-off, being either non-parametric–and thus incapable of generalizing to new data without re-training the whole model from the beginning–or linear–that leads to inaccuracy alignment.

Besides, for improving phenotype prediction, feature selection and/or extraction (unsupervised learning) are widely used as a preprocessing step prior to supervised learning. However, since the preprocessing step is separated from the predicting step, highly predictive features may have missed and thus affect the prediction performance. For instance, many disease genes are not differentially expressed between disease and control [3]. To address this, regularization is used as complementary approaches. Basically, regularization imposes prior information to the supervised learning models. For example, previous methods impose the L1 regularization for implicitly selecting features [10]. Other methods apply the Laplacian regularization for imposing feature networks such as gene regulatory networks and protein-protein interactions [9]. Instead of penalizing each network edge equally as in Laplacian regularization, another method penalizes each network feature equally [14]. However, these regularizations are from general biological knowledge, lacking the rich details of concrete input biological datasets at hand, especially the distance metric among all possible features of the inputs.

To address above issues, we developed an interpretable regularized learning model, deepManReg to predict phenotypes from multi-modal data (Fig. 1). In particular, deepManReg simultaneously (1) identifies nonlinear multi-modal relationships and (2) predicts phenotypes from multi-modal features and relationships. In particular, it first learns coupled deep neural networks to align cross-modal manifolds onto a common latent space. This step aims to preserve both global consistency and local smoothness across modalities and reveal higher-order nonlinear cross-modal relationships and, especially, solved the trade-off between nonlinear and parametric manifold alignment. Second, deepManReg uses cross-modal manifolds as a feature network [14] to regularize the classifier for improving phenotype predictions and also prioritizing the features and cross-modal interactions for the phenotypes. To solve this learning problem, we developed a novel optimization algorithm by backpropagating the Riemannian gradients on a Stiefel manifold. We applied deepManReg to recental single cell mutli-modal data such as transcriptomics and electrophysiology for neuronal cells in the mouse visual cortex. We show that deepManReg improves predicting cellular phenotypes (e.g., cellular layers) and also prioritizes genes and electrophysiological features for the phenotypes. Finally, deepManReg is open-source and general for phenotype prediction from multi-modal data.

**Figure 1:**
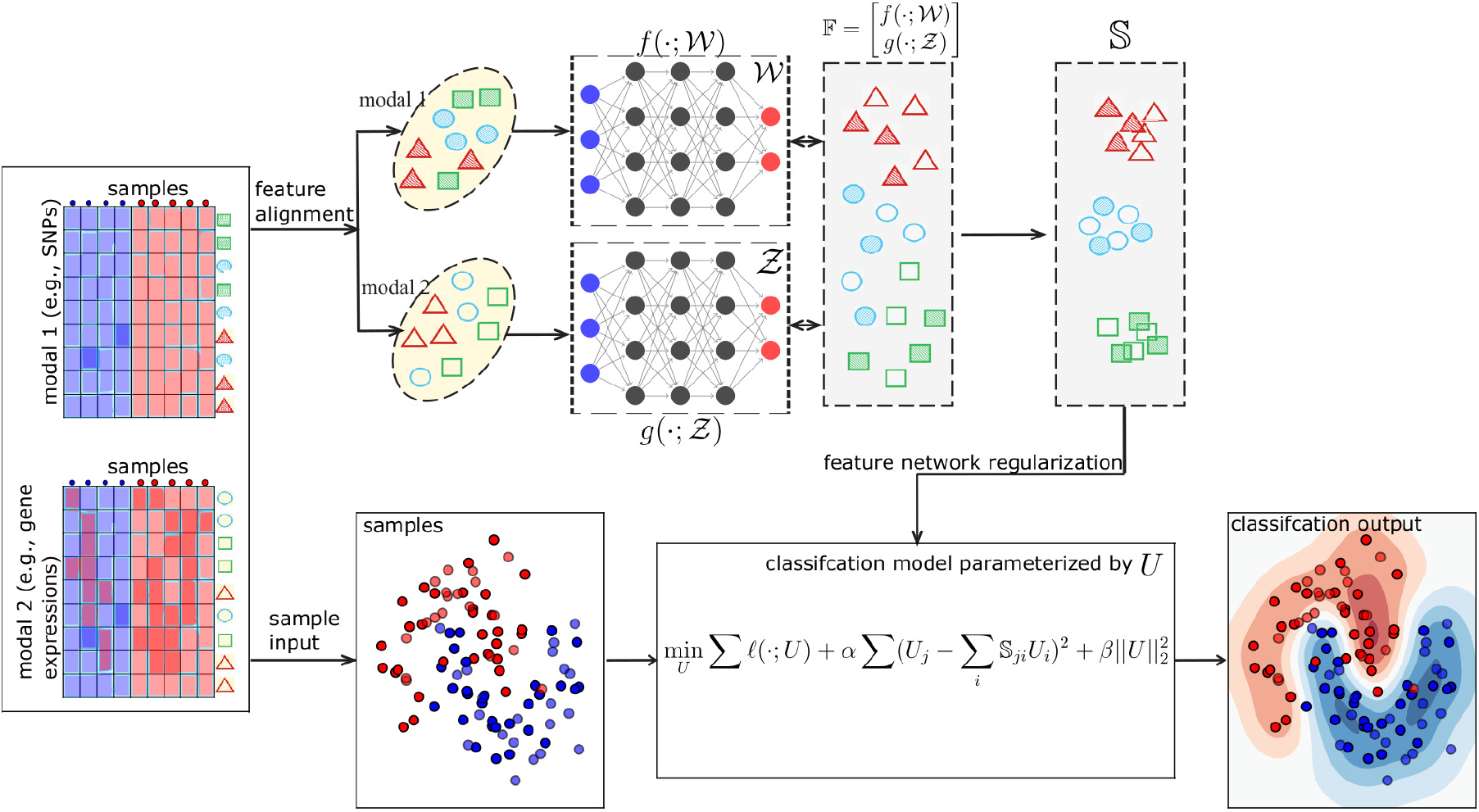
deepManReg: a deep manifold-regularized learning model for improving phenotype prediction from multi-modal data. deepManReg inputs multi-modal datasets, e.g., Modal 1 (left top) and Modal 2 (left bottom), across the same set of samples. In Phase 1 (top flow), deepManReg aligns all features (the rows) across modalities by deep manifold alignment. In particular, it uses coupled deep neural networks 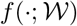 and 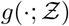, parameterized with 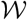 and 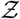 to project the features onto a common latent manifold space 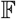. The similarity matrix 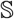 of features on the latent space is then calculated, encoding the similarity of nonlinear manifolds among all pairs of both cross-modal and within-modal features. In Phase 2 (bottom flow), deepManReg inputs all the samples (the columns of the input data) into a regularized classification model, parameterized by *U*. The similarity matrix of features on the latent manifold space 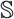 in Phase 1 is used to regularize this classification model (i.e., via feature graph regularization), imposing similar features to have similar weights, when training. Finally, deepManReg outputs a regularized classification (i.e., deep manifold-regularized) for improving phenotype prediction and prioritizing cross-modal features for the phenotypes.

Finally, It is also worthy noting that there are differences between deepManReg and other geometric-based learning methods: (a) structured-output learning methods, such as graph neural networks can only learn the structures or relationships among samples [16]; (b) graph-regularized learning methods (mostly based on Laplacian graphs) use the relationships of features to regularize the learning model but aim to penalize each network edge, rather than each feature itself [14].

## 2 Methods

Under our recent multiview learning framework [13], deepManReg inputs multi-modal data of samples, aligns multi-modal features and predicts the samples’ phenotypes. For instance, two modalities of a set of samples can be modeled as 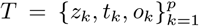 with 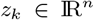 being the data of Modal 1 and 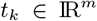 being the data of Modal 2, and associated phenotypes for both modalities (i.e., labels) 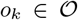. There are two major phases in deepManReg: (1) aligning multi-modal features by deep-neural-network based manifold alignment (deep manifold alignment) for identifying nonlinear, cross-modal feature relationships on a common latent space, and (2) predicting the phenotypes of the samples from both multi-modal data and cross-modal relationships via regularized classification.

### 2.1 Phase 1: Deep manifold alignment of multi-modal features

#### 2.1.1 Deep manifold alignment via parametric nonlinear alignment between the manifolds of multi-modal features

Manifold alignment is a class of techniques for learning representations of multiple data views, such that each view’s representation is simultaneously the most predictive of, and the most predictable by, the other. It can also be considered as a generalization of canonical correlation analysis (CCA) whereas the intrinsic geometry of data views are preserved and/or the projections are nonlinear [13].

Manifold alignment has been applied to identify linear (feature-level) projections, or nonlinear (instance-level) embeddings of multi-modal data. While the instance-level version generally produces more accurate alignments, it sacrifices a great degree of flexibility as the learned embedding is often difficult to parameterize. The feature-level projections allow any new instances to be easily embedded into a manifold space, and they may be combined to form direct mappings between the original data representations. These properties are crucial for transferring knowledge across modalities. Thus, deepManReg simultaneously learns different nonlinear mappings for different data modalities and align them onto a common manifold latent space. This idea combines appealing properties of both feature-level and instance-level projections for achieving accurate alignment and generalization. Furthermore, traditional solutions for manifold alignment rely on the eigendecompostion that is typically computationally intensive. To address this, we utilize stochastic gradient descent (SGD) as an implicit regularization and backpropagation for speeding up training deepManReg models.

Particularly, deepManReg first aims to calculate the similarities in terms of nonlinear manifolds among all possible features across modalities. To this end, deepManReg conducts a deep manifold alignment between all features so that the features are aligned onto a common latent manifold space. The distances of the features on the latent space thus reveal such similarities of the features for nonlinear manifold structures, suggesting nonlinear, cross-modal feature relationships. Mathematically, given two modal datasets, 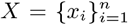 and 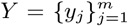 where 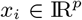 are the features of Modal 1 and 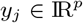 are the features of Modal 2, and the partial correspondences between the instances in *X* and *Y*, encoded by the matrix 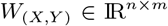, we want to learn the two mappings *f* and *g* that map *x_i_,y_j_* to 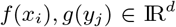 respectively onto the latent manifold space with dimension *d* ≪ *p* that preserves local geometry of *X*, *Y* and also matches cross-modal features from the correspondence.

Further, the instance *x_i_* is correspondent to the instance *y_j_* if and only if *f*(*x_i_*) = *g*(*y_j_*). Besides, any prior correspondence information between the features from different modalities can be used as partial information to initially build the corresponding matrix *W*(*X,Y*). After mappings, 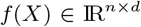 and 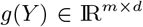 represents the new coordinates of the features of Modal 1 and Modal 2 on the latent manifold space with the dimension *d*, respectively. That said, the concatenation of the new coordinates 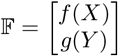 is the unified representation of the features from *X* and *Y* on the common latent manifold space.

Then, according to [21], the loss function for manifold alignment can be formed as the Laplacian eigenmaps [2] using the joint Laplacian and the joint adjacency matrix of the two datasets:

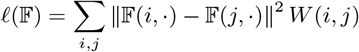

where the sum is taken over all pairs of instances from both datasets, 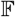 is the unified representation of both datasets, and 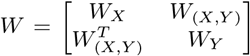 is the joint similarity matrix (*W_x_* and *W_Y_* are similarity matrices within each dataset *X* and *Y*).

Using the facts that 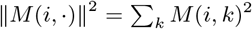 and that the Laplacian is a quadratic difference operator, the above equation can be transformed into:

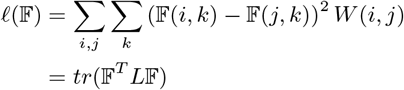

where *L* is the joint Laplacian [21] of both datasets.

For this loss function to work properly, i.e., avoiding the trivial solution of mapping all instances to zero, we need an additional constraint,

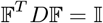

where *D* is the diagonal matrix of *W* and 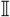 is the *d* × *d* identity matrix.

Then, we have this equation for manifold alignment:

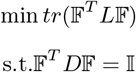

Normally, this optimization can be solved by eigendecomposition [22] which is computationally intensive. Moreover, solving generalized eigenvector problem gives us merely the new coordinates of the latent manifold (i.e., *X*’ = *f* (*X*), *Y*’ = *f* (*Y*)), not the closed form of mappings themselves (i.e., *f* (·) and *g*(·)), and thus is incapable of generalizing for new instances. To solve this, we parameterized the mappings *f* (·) and *g*(·) by using coupled deep neural networks and finally form the optimization problem as below:

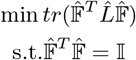

if we set 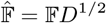 and 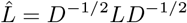.

This actually an optimization problem on the Stiefel manifold, where the feasible set of the orthogonality constraints 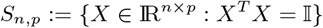 is referred to as the Stiefel manifold, which was due to Stiefel in 1935 [18].

#### 2.1.2 Deep Neural Networks to represent nonlinear embedding of manifold alignment

As above, we model the relationships between the observable data *x_i_, y_j_* and its latent representation *f* (*x_i_*),*g*(*y_j_*) using two nonlinear mappings 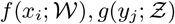 where 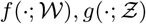 denote the mapping functions and 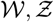 denote the set of the function parameters. In deepManReg, we employ the deep neural networks (DNNs) to model our mapping functions, since DNNs have the ability of approximating any continuous mapping using a reasonable number of parameters. Note that, of the two DNNs, the numbers of input features are unnecessary to be the same, but the numbers of output represented features have to be exactly the same for allowing having a common latent space. Precisely, if 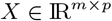 is a matrix of data vectors 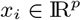, the number of input features for the first network 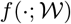, and if 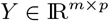 is a matrix of data vectors 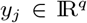, the number of input features for the second network 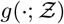 is *q*. The numbers of output represented features of both DNNs is *d*, the dimension of the common latent manifold space.

#### 2.1.3 Optimization on Stiefel Manifold to train two deep neural networks for parametric nonlinear manifold alignment problem

There exist two key issues for generalizing backprop to the context of training our DNNs for deep manifold alignment. The first one is preserving the manifold constraint in the output layer. As we force the outputs to be on Stiefel manifolds, merely using the forward propagation in the normal DNN cannot yield valid orthogonal outputs. While the gradient of loss function with respect to output layer, i.e., 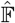, can be calculated easily, computing those with hidden layers, i.e. 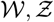 has not been well-solved by the traditional backprop, which may be the second key issue for training the DNNs in deepManReg.

**Algorithm 1:**
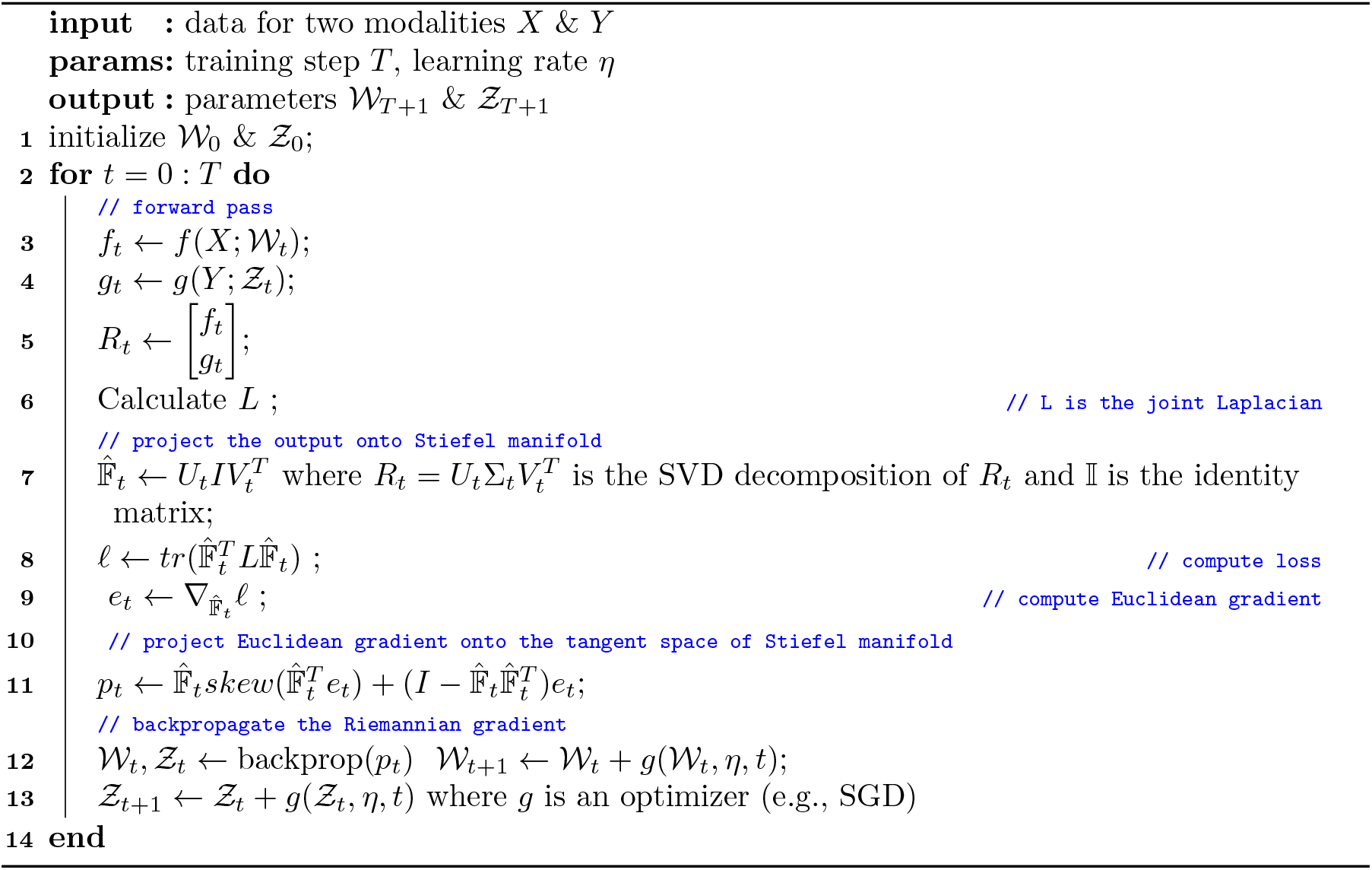
Deep Manifold Alignment

To solve the first issue of preserving the constraint, we construct the last layer by projecting the output of the preceding layer 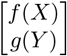 onto the Stiefel manifold *S_m+n,d_*. Specifically, we use the classical projection operator π(·) which is defined as:

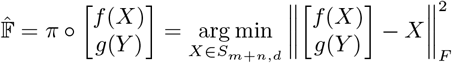

it is known that the solution of this problem is given by

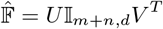

where 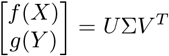 is the SVD decompostion of 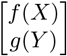. Thus, 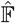 now is orthogonal output, i.e. 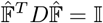

As for the second issue, we developed a new way of updating the weights 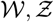 by exploiting an SGD setting on the Stiefel manifolds. The steepest descent direction for the corresponding loss function *ℓ(R)* with respect to *R* on the Stiefel manifold is the Riemannian gradient 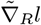 To obtain it, the Euclidean gradient 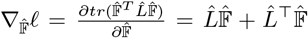 is projected onto tangent space 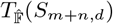 of Stiefel manifold *S_m,n,d_*. The projection is defined as

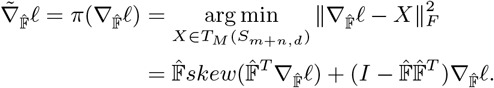

where *skew*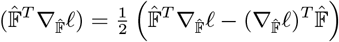.

Putting all together, we summarized our optimization in Algorithm 1, which can be readily implemented with the modern tools for automatic differentiation such as PyTorch.

### 2.2 Phase 2: Regularized classification by cross-modal feature relationships learned from deep manifold alignment

After finding the common latent space from deep manifold alignment, we can now calculate the distance matrix 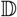 for each row pairs of matrix 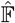, and then similarity matrix 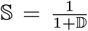. The latter finally gives the similarities of all multi-modal features in terms of nonlinear manifold structures, systematically revealing cross-modal feature relationships.

In Phase 2 of deepManReg, we want to improve phenotype prediction from multi-modal data using such cross-modal feature relationships. In particular, back to the training set 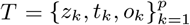, deepManReg learns a classifier paramaterized by a weight *U* by minimizing a loss function *ℓ*(*z,t,o; U*) over the training instances (*z_k_,t_k_,o_k_*) [14]. Now, with the similarity information of features, provided by matrix 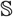 from the previous step, we can use 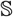 as an adjacency matrix of a feature graph encoding the relationship between all pairs of features within and across modalities. The degree of each vertex in the feature graph has to be sum to one, 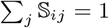, to avoid some features dominating the whole graph. Because similar features should have similar weights after training, we regularize each feature’s weight by the squared amount it differs from the weighted average of its neighbors. Thus, the loss function for feature network regularized learning is [14]:

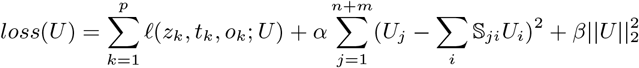

The hyperparameters *α* and *β* are to balance between the network regularization and the ridge regularization. Finally, the combined regularization can be rewritten as *U^T^MU* where 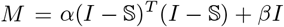.

The classifiers can be general. In practice, here, we use a neural network as a classifier so the optimization problem above can be solved easily with gradient descent methods. Also, we can use other approaches for regularization such as the graph Laplacian [14]. The main difference between Laplacian regularization and feature graph regularization is that Laplacian penalizes each edge (between two features) equally while the latter penalizes each feature (e.g., nodes) equally. The efficiency of the approaches should depend on the problem domain.

## 3 Results

### 3.1 Predicting cellular phenotypes from single cell multi-modal data

Recent Patch-seq technique measures multi-modal characteristics of single cells such as transcriptomics, electrophysiology and morphology [4]. For example, the Brain Initiative project has generated multi-modal data of neuronal cells in the human and mouse brains [6]. Using those single-cell multi-modal data, ones have identified many cell types corresponding to various cellular phenotypes. Here, we applied deepManReg to recent Patch-seq data for the mouse visual cortex from Allen Brain Atlas for predicting neuronal phenotypes, including cell layers and transcriptomic types. Specifically, this dataset includes the transcriptomic, and electrophysiological data of 4435 neuronal cells (GABAergic cortical neurons) in the mouse visual cortex [6]. For cellular phenotypes for our prediction, we included six major transcriptomic cell types (t-types): Vip, Sst, Sncg, Serpinf1, Pvalb, and Lamp5, and five cell layers revealing the locations of cells on the visual cortex: L1, L2/3, L4, L5, and L6.

### 3.2 Datasets and data preprocessing

The electrophysiological data includes the responses of three stimuli: short (3 ms) current pulses, long (1 s) current steps, and slow (25 pA/s) current ramp current injections. We extracted 47 electrophysiological features (e-features) on stimuli and responses, identified by Allen Software Development Kit (Allen SDK) and IPFX Python package [1]. We then filtered the e-features with many missing values, extracted the cells from t-types and layers as above, and finally selected 41 e-features for 3654 neuronal cells. The transcriptomic data quantifies gene expression levels of the neuronal cells on the genome wide. In particular, we extracted the 500 genes that have the highest express variations among the 3654 cells. Then, we input the gene expression and e-features of those cells as input multi-modal data into deepManReg for predicting cellular phenotypes, i.e., *X* is 500 genes by 3654 cells and *Y* is 41 e-features by 3654 cells. As shown in Fig. 2, the latent space from deep manifold alignment (Phase 1) reveals that many genes and e-features have strong nonlinear relationships (via aligned cross-modal manifolds).

**Figure 2:**
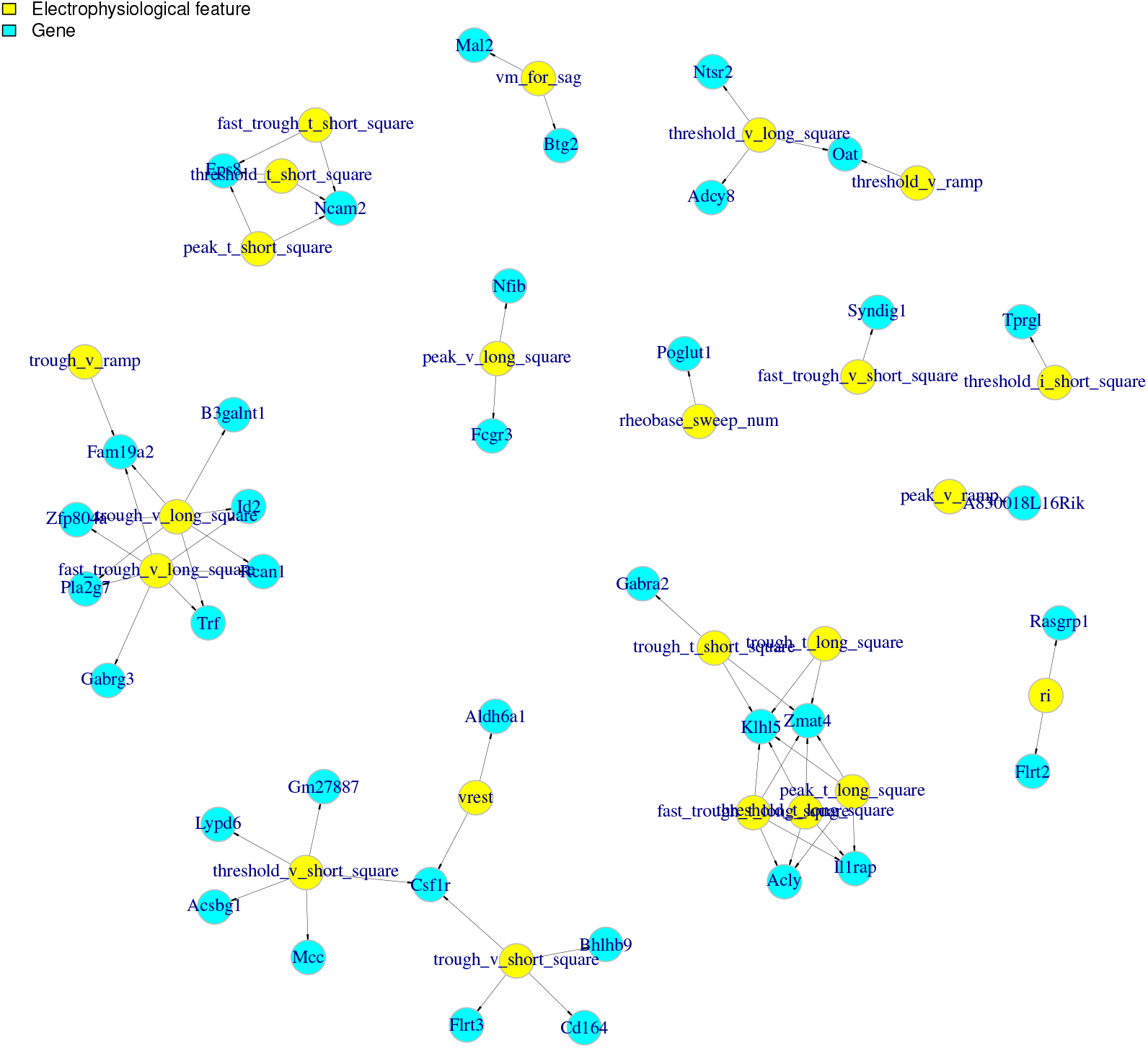
The genes and electrophysiological features (e-features) of neuronal cells in the mouse visual cortex having highly correlations of their reduced dimensions on the aligned latent space by deepManReg Phase 1 (i.e., deep manifold alignment). Cyan: genes. Yellow: e-features. Nodes are connected by correlations of reduced dimensions (10-dim) > 0.85.

### 3.3 deepManReg improves cellular phenotype prediction from single cell transcripotimics and electrophysiology

After deep manifold alignment, we applied deepManReg to use the aligned latent spaces of genes and e-features to regularize another deep neural network model to classify the cellular phenotypes such as cell layers. In particular, the neural network for classification has the input layer consisting of 541 nodes (500 genes + 41 e-features), two hidden layers (100/50 hidden units) and the final output layer with the same number of units as phenotypes along with a Softmax operation. For instance, for classifying cell layers, the output units represent L1, L2/3, L4, L5, and L6. We randomly split all cells into the training/testing sets with a stratified ratio of 80/20 and obtained 500 sets. For each training set, we oversampled the cells from each label to be 941 cells and thus balance sample sizes across labels (e.g., L1: 262 cells; L2/3 1097 cells; L4: 385 cells; L5: 1176 cells; L6:734 cells) [15]. As shown in Fig. 3A, the prediction accuracy of deepManReg for the testing sets to classify cell layers is significantly higher than the classification without any regularization (k.s. test p¡ 7.85*1e-13). Also, its average accuracy, 44.6% (with a 95% confidence interval [28.0%, 53.0%]) is higher than both the baseline of 20% (five labels) and the average accuracy of the classification without regularization (39.7% mean accuracy, [7.1%, 52.8%] confidence interval). Moreover, as shown in Fig. 3B, deepManReg also achieves relatively high AUC values of 0.93, 0.81, 0.74, 0.72, and 0.87 for the five layers L1, L2/3, L4, L5, and L6, respectively. In addition to predicting cell types, we also found that deepManReg outperforms the classification without regularization for predicting t-types (k.s., test p¡ 1.62*1e-9), i.e., average accuracy 83.2% ([62.5%, 93.3%] confidence interval) for deepManReg vs. 80.8% ([34.1%, 95.9%] confidence interval) for the latter. These results demonstrate that the regularized classification by deep manifold alignment improves predicting cellular phenotypes from single cell multi-modal data. This also suggests potential contributions from the nonlinear manifold relationships of gene expression and electrophysiology to the cellular phenotypes.

**Figure 3:**
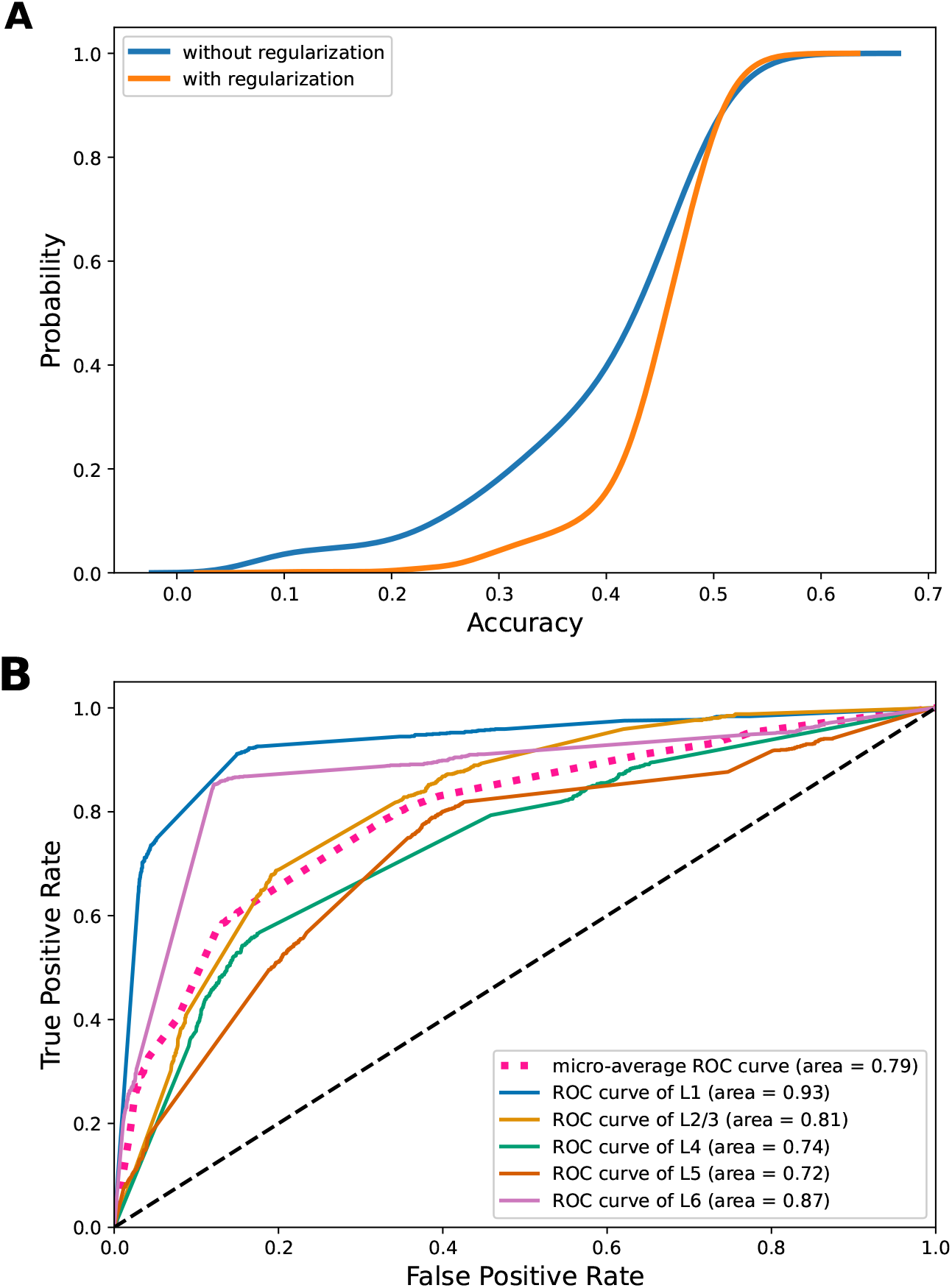
(A) Cumulative distributions of testing accuracies for classifying cell layers in the mouse visual cortex by deepManReg (Blue) vs. the neural network classification without any regularization (Orange). (B) Receiver operating characteristic (ROC) curves for classifying cell layers in the mouse visual cortex by deepManReg. Blue: L1, Yellow: L2/3, Green: L4, Orange: L5, Purple: L6. x-axis: False Positive Rate, y-axis: True Positive Rate.

### 3.4 Prioritization of multi-modal features for cellular phenotypes via integrated gradients

After training a deepManReg model, we further used a derivative-based method called integrated gradient [20] for prioritizing genes and e-features for each phenotype (e.g., cell layers in Supplemental Table 1). Specifically, we calculated the gradient of the model’s prediction for each e-feature and/or gene to quantify the changes of the output response values (e.g., cell layers) by a small change of input gene expression and e-feature values [12]. We used the recent Python package, Captum [7] to implement the integrated gradient method and calculate the importance scores of each gene/e-feature for output labels (i.e., cellular phenotypes). We then ranked the genes and e-features by the scores and prioritized top ones for each phenotype. For instance, we summarized top prioritized genes and e-features for each cell layer in Supplement Table 1.

## 4 Conclusion

In this paper, we presented a novel regularized learning method, deepManReg for simultaneously (1) revealing nonlinear manifold relationships across multi-modal data and (2) improving phenotype predictions via regularization by cross-modal manifolds. In particular, deepManReg learns multiple deep neural networks for different modalities and jointly trains them to align multi-modal features onto a common latent space. The distances of various features in and between modalities on the space represent their nonlinear relationships identified by cross-modal manifolds.

Although we demonstrated that deepManReg works for two particular datasets (i.e., brain disorders and single cells), deepManReg can be generalized to any multi-modal data such as additional single cell omics (scATAC-seq, scHi-C, etc). Also, its deep neural networks for manifold alignment can be designed specific for each modality. For example, if two modalities are genomics and images, the neural network for aligning images can be changed to a convolutional neural network. Also, one can model those neural networks by recent graph neural networks [16], aiming to not only align multi-modal features but also underlying biological networks in the modalities.

deepManReg solves the tradeoff between nonlinear and parametric manifold alignment (by utilizing the nonlinearity and parametric of neural architecture which is trained by a Riemannian optimization procedure). Besides, deepManReg works as both representation learning and regularized classification. However, training Deepalignomic requires a non-trivial hyperparameter optimization since training two deep neural networks simultaneously includes a large combination of parameters. Another potential issue for aligning such large datasets in deepManReg which may be computational intensive is the large joint Laplacian matrix (Algorithm 1). Therefore, in future, we may use the Nystrom method [5] to approximate the Laplacian matrix for making deepManReg more scalable.

## Funding

This work was supported by the grants of National Institutes of Health, R01AG067025, R21CA237955 and U01MH116492 to Daifeng Wang and U54HD090256 to Waisman Center.

## Conflict of Interest

none declared.

## Notes

### Competing Interest Statement

The authors have declared no competing interest.

